# Whole-brain functional connectivity neuromarkers uncover the cognitive recovery scheme for overt hepatic encephalopathy after liver transplantation

**DOI:** 10.1101/2020.09.01.278614

**Authors:** Yue Cheng, Wen Shen, Junhai Xu, Rachel C. Amey, Li-Xiang Huang, Xiao-Dong Zhang, Jing-Li Li, Cameron Akhavan, Ben A. Duffy, Wenjuan Jiang, Mengting Liu, Hosung Kim

## Abstract

Neurocognitive impairment is present in cirrhosis and may be more severe in cirrhosis with the overt hepatic encephalopathy (OHE). Liver transplantation (LT) may reverse the impaired brain function. MRI of resting-state functional connectivity can help unravel the underlying mechanisms that lead to these cognitive deficits and recovery. Sixty-four cirrhotic patients (28 with OHE; 36 without) and 32 healthy controls were recruited for resting-state fMRI. The patients were scanned before and after LT. We evaluated pre- and postsurgical neurocognitive performance in cirrhotic patients using psychomotor tests, i.e. number connection test (NCT) and digit symbol test (DST). Network-based statistics found significant disrupted connectivity in both groups of cirrhosis with OHE and without compared to controls. However, the presurgical connectivity disruption in patients with OHE was included in a greater number of connections than those without (65 vs. 17). The decrease in FC for both OHE and non-OHE patient groups was reversed to the level of controls after LT. An additional hyperconnected network (i.e., higher than controls) was observed in OHE patients after LT (p=0.009). Regarding the neural-behavior relationship, the functional network that predicted cognitive performance in healthy individuals, showed no correlation in presurgical cirrhotic patients. Such an impaired neural-behavior relationship was re-established after LT for non-OHE patients but not for OHE. OHE patients displayed abnormal hyperconnectivity and persistently impaired neural-behavior relationship after LT. Our results suggest that patients with OHE may undergo a different trajectory of postsurgical neurofunctional recovery in comparison to those without, which needs further clarification in the future study.

## Introduction

Overt hepatic encephalopathy (OHE) is one of the most prominent complications in cirrhotic patients. It decreases a patient’s quality of life in addition to lowering their survival rate. Neurocognitive impairment is present in cirrhosis and occurs along a continuous spectrum. These impairments are accelerated by episodes of OHE (Bajaj et al., 2010), which can often irreversibly impair attention, learning ability, and executive function (Campagna et al., 2014). Studies have demonstrated (Bajaj et al., 2010; Umapathy, Dhiman, Grover, Duseja, & Chawla, 2014) poorer neurocognitive performance in patients with OHE than those without.

Liver transplantation (LT) is the only effective treatment for end-stage cirrhosis (Cárdenas & Ginès, 2011). Successful LT can restore liver function completely, however, to what extent cognitive dysfunction can be recovered, and to what extent OHE can impact the recovery process are still unclear (Ahluwalia et al., 2016; Cheng et al., 2017; Sotil, Gottstein, Ayala, Randolph, & Blei, 2009).

Neurocognitive deficits and recovery are usually assessed by extensive psychometric neurocognitive testing for cirrhosis studies. So far there is clear evidence suggesting that multiple cognitive functions are impaired by cirrhosis (Campagna et al., 2014). Functional MRI, particularly resting-state functional connectivity (Chen et al., 2016; Cheng et al., 2017; Qi et al., 2012; Qi et al., 2014; Zhang, Cheng, Liu, & behavior, 2017; Zhang, Cheng, Shen, et al., 2017), could help further unravel the underlying mechanisms that lead to these cognitive deficits and recovery. Several studies have investigated the association between the impaired cognitive functions and the functional connectivity related to these functions using hypothesis-driven approaches (García-García et al., 2018; Qi et al., 2012). However, which localized functional connectivity corresponds to which cognitions is still not well established yet for many cognitive functions (Barch et al., 2013; R. A. Poldrack & Yarkoni, 2016; R. A. J. P. o. P. S. Poldrack, 2010). Furthermore, studies of patients with various brain injuries have evidenced that functional hyperconnectivity or brain reorganization may occur in response to the initial brain injury, leading to broader connectivity changes than the changes in the relavent brain regions (Bernier et al., 2017; Bharath et al., 2015; Castellanos et al., 2010; Grefkes & Ward, 2014). This impedes the investigators to seek a very clear conclusion on their neuroimaging data, when applying hypothesis-driven approahes where only hypothetically relavent functional regions are investigated. Hence, searching the possible biomarkers across the whole brain network, via data-driven approaches, without restricting within known brain functional regions, connections, and functional networks would be critical to clarify the cirrhosis associated neurocognitive impairments and recovery.

In this study, we used a data-driven approache that combined cognitive task with whole-brain FCs to provide a new perspective on cognitive deficits and recovery for OHE and no-OHE cirrhotic patients before and after the transplant surgery. We evaluated neurocognitive deficits in cirrhotic patients and recovery after LT using a standard psychomotor cognitive task that is reliant on a variety of functional neural components like sensorimotor, attention, memory, and even executive functions (Miyake et al., 2000). Without restricting our study in pre-hypothesized brain regions, connections, and functional networks, we searched the possible biomarkers across the whole brain. Based on the functional connectivity found using data-driven approaches, we wanted to address questions such as: ***1)*** are the disrupted or altered functional connections in cirrhotic patients with OHE or those without?; ***2)*** if so, are these connecitons recovered, reversed or reorganized after LT treatment? ***3)*** Then, are the re-established connections after LT the same as controls or different from controls due to brain reorganization? ***4)*** Is the normal relationship observed between healthy brain and cognitive performance impaired in presurgical cirrhotic patients and is this recovered after the LT?

## Methods

### Subjects

In this prospective study, approval was obtained from the Ethics Committee of our institutional review board. All subjects provided the written informed consent before being included in the study. From November of 2013 to January of 2018, 64 patients with end-stage cirrhosis (28 Patients with the history of OHE episodes, and 36 Patients without) scheduled to undergo LT were recruited from the department of Transplantation Surgery. 32 of the 64 patients have been previously reported (Cheng et al., 2015; Zhang, Cheng, Shen, et al., 2017) investigating the regional functional activity, whereas in this study we evaluated the functional connectivity and specifically its predictive power to behaviors. LT candidates who completed all necessary laboratory examinations, neuropsychological tests, and baseline MRI were included. The exclusion criteria were the following. Participants were excluded if they reported: (1) a history of drug or alcohol abuse; (2) presence of any noticeable brain lesions on conventional MR, such as a tumor or stroke; (3) any major neurologic or psychiatric disorders; (4) history of liver cancer; (5) previous liver or other organ transplantation; (6) head motion of 1.5 mm or 1.5° during MRI. The etiology of the cirrhotic patients included type B hepatitis (n = 35), type C hepatitis (n = 15), primary biliary cirrhosis (n = 8), and cryptogenic cirrhosis (n = 6). Thirty-six of these patients eventually received successful LT and completed the one-month follow-up examination. Included patients had no complications such as acute transplant rejection, liver failure, severe biliary complications, or any neurologic complications, such as alterations in mental status, seizures, and focal motor deficits. A detailed flowchart of this study is provided in Figure 1.

**Figure 1.**
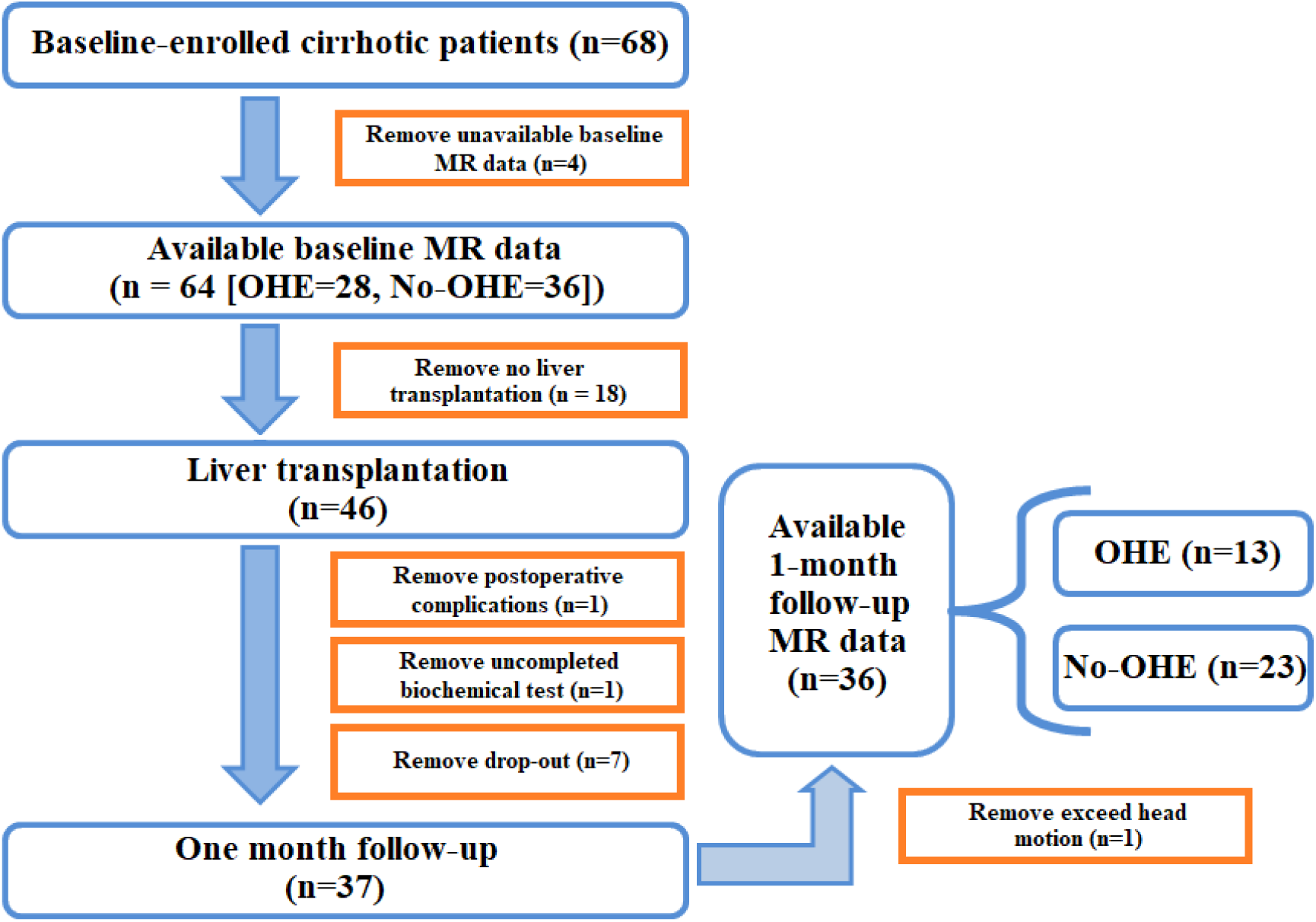
Subjects involved in this study. 68 cirrhotic patients were enrolled in this study, part of them were removed from the subject pool for the following reasons: unavailable baseline MR data, not undergoing LT, postoperative complications, incomplete biomedical exam, dropping out, and having excess head movement (see panels for exact numbers). In total, 28 patients with OHE and 36 without OHE were analyzed before LT; 13 patients with history of OHE and 23 patients without history of OHE participate in the study after the LT operation.

Thirty-two age- and sex-matched healthy controls (HCs) were recruited from the local community. All HCs had no diseases or history of liver or neurologic diseases. All HCs were self-identified as right-handed with normal sight and had completed neuropsychological tests (see details in *Neuropsychological tests*).

### Laboratory examinations

All patients completed blood laboratory tests to evaluate liver functioning one week prior to their MRI scan both before and one month after LT. These tests included prothrombin time, albumin, and total bilirubin. For the preoperative patients, their Child-Pugh score was used to assess liver dysfunction. Venous blood ammonia was also tested for cirrhotic patients. No blood laboratory tests were performed for HCs.

### Neuropsychological tests

All patients and HCs underwent two typical neuropsychological tests to evaluate cognitive function before the MRI scan (collected by Y.C. and W.S.). These tests included the number connection test type A (NCT-A; are abbreviated as NCT in all the texts below) and digit symbol test (DST) (Karin Weissenborn, Ennen, Schomerus, Rückert, & Hecker, 2001). The NCT and DST are parts of the Psychometric Hepatic Encephalopathy Score (Ferenci et al., 2002; K. Weissenborn et al., 2005), and are ideally suited as they are supported by a variety of functional neural components given its reliance on different sensorimotor, attention, memory, and even executive functions (Miyake et al., 2000). In NCT task, participants were required to connect randomly placed figures in order as quickly as possible to measure psychomotor speed and attention (Bajaj, Wade, & Sanyal, 2009). Completion time is indicative of performance, and worse performance is indicated by a longer completion time. The DST is a measure of complex visuomotor tracking and learning, which is relatively unaffected by intellectual ability, memory, or learning. This task emphasizes sustained attention, response speed, and visuomotor integration (Bajaj et al., 2011). Digits from one to nine and corresponding symbols are displayed in front of the subjects, who are then asked to fill in the blanks with the symbol that matched each digit. Participants have 90 seconds to complete this task. The number of correctly transcribed symbols was used as an indicator of performance - lower scores indicated underperformance.

### MR Imaging Data Acquisition

A 3-T MR imager (TIM-Trio, Siemens Medical Solutions, Erlangen, Germany) was used to collect the imaging data (collected by Y.C. and X.D.Z). For functional images, blood oxygen level-dependent single-shot echo-planar sequence (repetition time/echo time, 2500/30 ms; flip angle, 90°; field of view 220 ×220 mm2; matrix, 96 ×96; iPAT, 2; number of slices, 40; slice thickness, 3 mm; intersection gap, 0.3 mm; 200 volumes; acquisition time, 8.5 minutes) was performed. During the scan, all subjects were instructed to close their eyes and remain awake, they were asked to think of nothing in particular. For structural images, sagittal 3D T1-weighted magnetization-prepared rapid acquisition gradient echo (MPRAGE) sequence was used with the following settings: 1900/3; inversion time (ms), 900; flip angle, 9°; number of slices, 176; slice thickness; 1 mm; matrix, 256 × 256.

### Data Preprocessing

Functional imaging data was preprocessed by J.X and R.C.A (MD and neuroscientist) using SPM12 software (https://www.fil.ion.ucl.ac.uk/spm/). For each participant, the first ten volumes were removed to allow for dynamic equilibrium and adaption to the scanning circumstances. The remaining functional images were corrected for time delays between slices using slice timing. A six-parameter rigid body transformation in the realignment analysis was performed to correct the head motion. Participant with a translation exceeding 3 mm and a rotation greater than 1.5 degree were not included in the analyses. In the present study no participants were removed. The individual subject’s structural image was then segmented into the gray matter, white matter, and cerebrospinal (CSF) for normalization after co-registration to the mean functional image. All functional images were spatially normalized to the standard Montreal Neurological Institute (MNI) space using the generated parameters at an isotropic voxel size of 3 mm. Finally, normalized functional images were smoothed with a Gaussian kernel (FWHM = 4 mm). A linear detrended and temporally band-pass filtered (0.01Hz < f < 0.08 Hz) procedure was performed to reduce the effects of low-frequency drift and high-frequency physiological noises (Biswal, Yetkin, Haughton, & Hyde, 1995). A linear regression was then used to reduce physiological noise and remove artifacts with several sources of spurious variance, including averaged signals from WM, CSF, and six head motion parameters.

### Whole-brain functional connectivity analysis

To obtain the whole-brain functional connectivity (FC) matrix, a prior Anatomical Automatic Labeling template (AAL) was used to divide the whole brain into 116 anatomical regions of interest (ROIs), including 78 cortical, 12 subcortical and 26 cerebellar regions (Tzourio-Mazoyer et al., 2002). A representative time series was extracted by averaging the time series of all voxels within each ROI. Then, Pearson’s correlation analyses were performed between each pair of ROIs to calculate the correlation coefficients, followed by the normalization with a Fisher z-score transformation. A symmetric functional connectivity matrix (116 × 116) was generated for each subject. The triangular portion of the adjacency matrix was extracted and transformed to a vectoral feature space with 6670 dimensions.

### Network based statistic (NBS)

To localize specific pairs of brain regions between which functional connectivity was altered in cirrhotic patients, we used the network based statistic (NBS) approach (Zalesky, Fornito, & Bullmore, 2010). Briefly, we first performed two-sample one-tailed t-tests in an element-by-element manner on those connections that were significantly non-zero (p < 0.05, Bonferroni-corrected) in at least one participant. Then, a primary threshold (p < 1 × 10^−4^ in this study; Wang et al., 2013) was applied to define a set of suprathreshold links within which any connected components and their size (defined as the number of links included in these components) were determined. To estimate the significance for each component, a null distribution of connected component size was derived empirically using a nonparametric permutation approach (10,000 permutations). For each permutation, all subjects were reallocated randomly into two groups, and two-sample one-tailed t-tests were conducted for the same set of connections mentioned above. The same primary threshold ((p < 1 × 10^−4^) was then used to generate suprathreshold links within which the maximal connected component size was recorded. Finally, for a connected component of size M found in the right grouping of controls and patients, the corrected p-value was determined by calculating the proportion of the 10,000 permutations for which the maximal connected component was larger than M.

### Connectome-behavior predictive mapping (CPM)

Connectome-behavior predictive modeling and all statistics were assessed by M.L. To investigate the reliable biomarkers in predicting task performance in resting-state whole brain FCs, a completely data driven approach of CPM was utilized for HCs. CPM searched all possible pairs of regions and their associated connectivity values, constructing a model (the “connectome”) that maximally fitted behavioral scores (Finn et al., 2015; M. Rosenberg, Finn, Scheinost, Constable, & Chun, 2017; Shen et al., 2017) using cross-validation. Specifically, linear regressions were run between each edge in the connectivity matrix in the resting state and DST/NCT scores. The resulting P-value for each regression was recorded in a 116 × 116 symmetric matrix, amounting to 6670 different linear regression significance values. To find the most meaningful associations between specific connectivity and the task performance, the resulting r values were held to a statistical threshold of p < 0.01 and separated into positive (edges whose connectivity strength indexed higher performance scores across subjects) and negative tails (edges whose connectivity strength indexed lower performance scores across subjects).

A single summary statistic, network strength, was used to characterize each participant’s degree of connectivity in the positive and negative tails. Positive network strength was calculated by averaging the edge strengths (Fisher-normalized r values) from a participant’s connectivity matrix in the edges of the positive tail, and negative network strength was calculated by summing the r values of the edges in the negative tail. Finally, network strength was utilized to predict performance scores across subjects using cross validation (Liu et al., 2020; M. D. Rosenberg et al., 2016).

Meaningful connectivity-based neuromarkers discovered from CPM in controls were then applied in cirrhotic patients before and after LT as a post-hoc analysis to evaluate whether the network can still predict subjects’ task performance, which subsequently evaluate the recovery of brain-behavior relations (Figure 4C).

**Figure 2.**
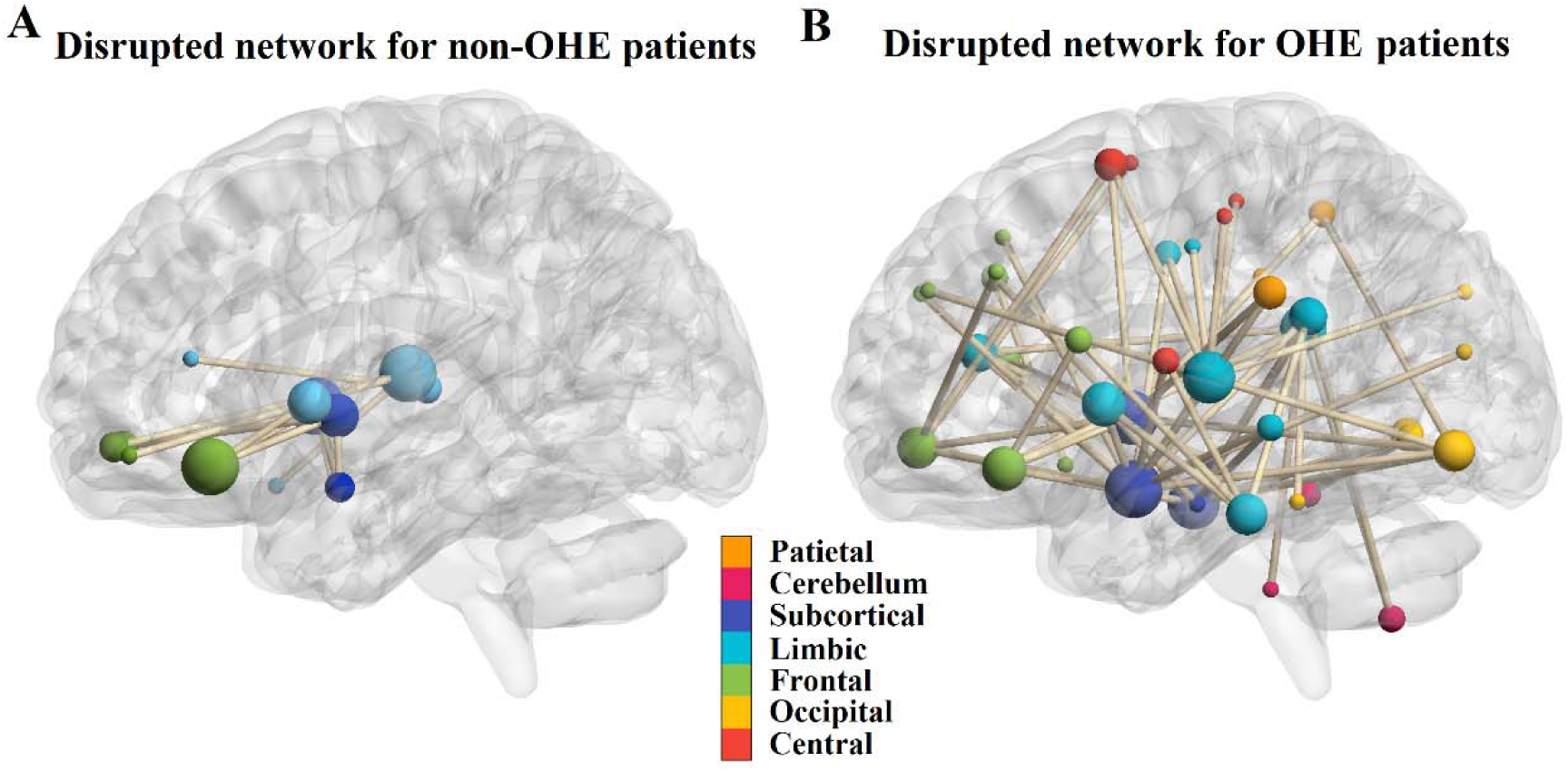
The disrupted resting-state functional networks for pre-LT non-OHE and OHE patients detected by NBS. (A) The region pairs showing significantly decreased functional connections in non-OHE patients compared to controls. These connections formed a single connected network with 16 nodes and 17 connections, which was significantly (NBS, p = .037, corrected) abnormal in the patients. The decreased connectivity was mostly found in the connections among sub-cortical nuclei such as Amygdala, Putamen, Pallidum and orbital fontal cortex. (B) The region pairs showing decreased functional connections in OHE patients. These connections formed a single connected network with 47 nodes and 65 connections, which was significantly (NBS, p = .006, corrected) abnormal in the patients. The connectivity was mostly found in the connections involving the following structures: sub-cortical nuclei such (Amygdala, Putamen, Pallidum), limbic system (temporal cortex, cingulate cortex, parahippocampal gyrus), orbital frontal cortex and other frontal, parietal cortex and cerebellum brain regions. OHE=overt overt hepatic encephalopathy, non-OHE = none-overt hepatic encephalopathy.

**Figure 3.**
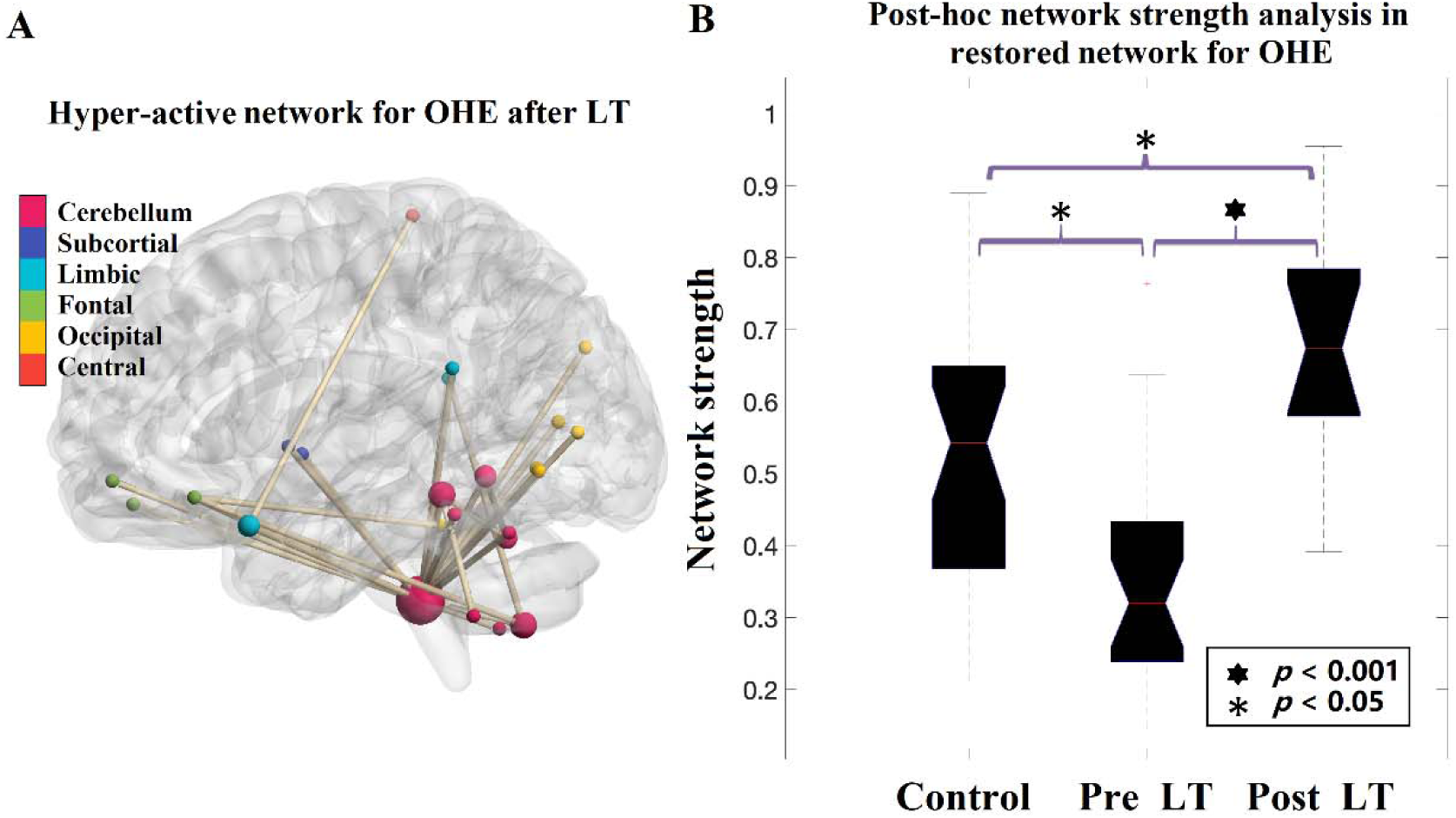
The restored resting-state functional network for post-LT OHE patients detected by NBS. (A) The region pairs showing increased functional connections in OHE patients after LT compared to pre-LT scans. These connections formed a single connected network with 19 nodes and 17 connections, which was significantly (p = .009, corrected) higher after LT. The increased connectivity was mostly found in the connections between cerebellum sub-regions, and between cerebellum and occipital cortex, orbital-frontal cortex and limbic systems. (B) Post-hoc analysis comparing the strength of the network detected in post-LT OHE patients in three groups. The strength of the network after LT was significantly higher than controls (two-sample t-test, p = 0.008, corrected), suggesting a LT-induced hyperconnectivity within the detected network. (C) Post-hoc analysis of comparing every single connectivity within the network between post-LT OHE patients with controls. OHE=overt hepatic encephalopathy, LT=liver transplantation.

**Figure 4.**
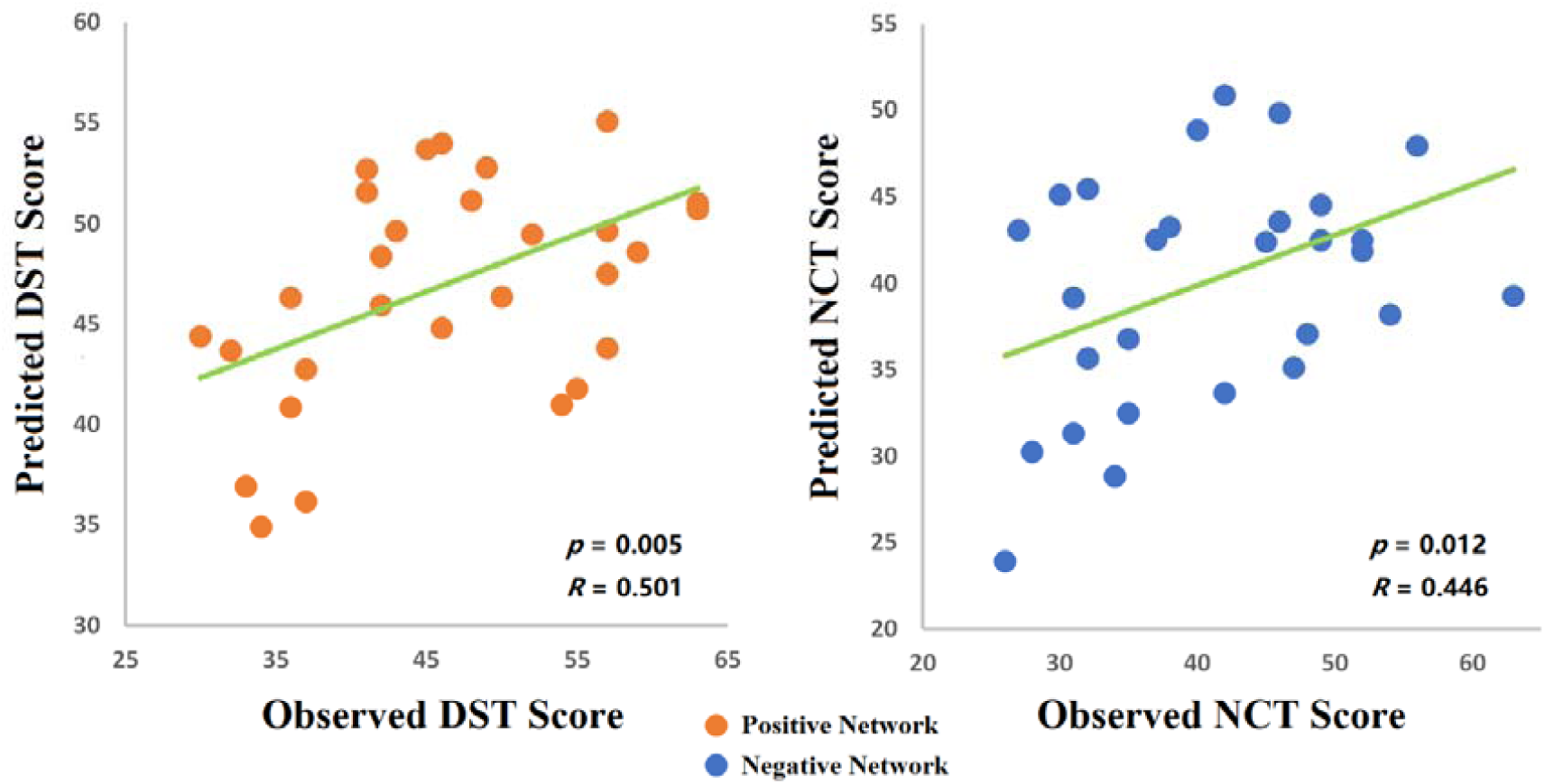
Functional connectivity models predicting cognitive task performance in healthy individuals. Scatter plots show correlations between observed performance scores and predictions by positive (left: for DST) and negative (right: for NCT) networks. Network models were iteratively trained on resting-state data from n−1 subjects in the control group and tested on resting-state data from the left-out individual. For negative networks in DST task and positive networks in NCT network, no meaningful correlations were found.

### Data availability

Anonymized data will be shared by request from any qualified investigator who provides a methodologically sound proposal, or for the purpose of replicating procedures and results presented in present study.

## Results

### 1. Demographics and Clinical Data

Demographic and clinical data for all subjects are summarized in Table 1. There were no meaningful differences in sex, age, or education level between OHE, no-OHE, and HC groups (two-way t-test, p-values > 0.05). For both no-OHE and OHE group, the liver function improved significantly (albumin, total bilirubin, blood ammonia, p≤0.01) or showed a tendency of restoration (prothrombin time) one month after LT (Table 1).

**Table 1.**
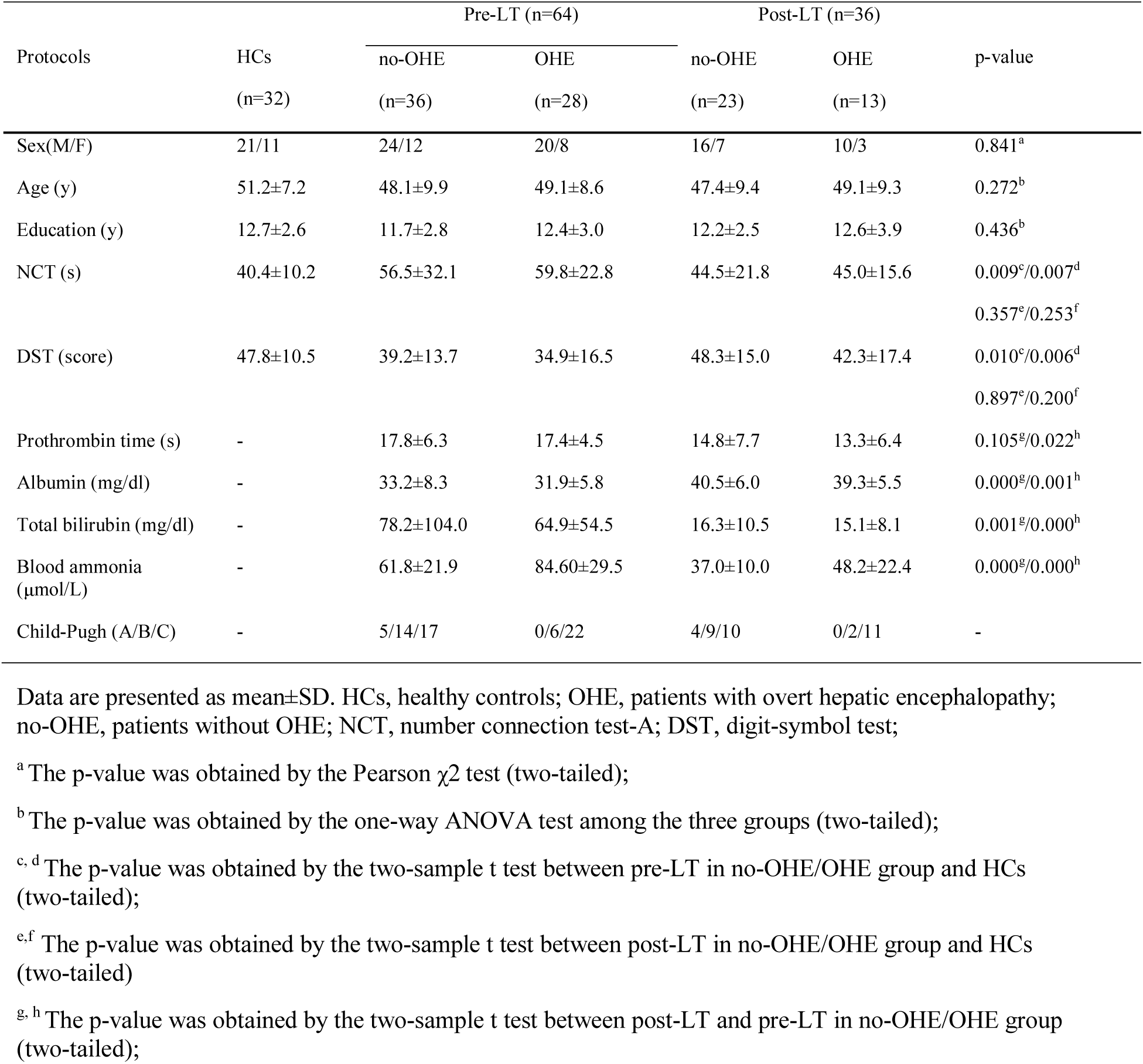
Demographic, neuropsychological, and biochemical information from the dataset.

| Protocols | HCs<br>(n=32) | Pre-LT (n=64) |  | Post-LT (n=36) |  | p-value |
| --- | --- | --- | --- | --- | --- | --- |
|  |  | no-OHE<br>(n=36) | OHE<br>(n=28) | no-OHE<br>(n=23) | OHE<br>(n=13) |  |
| Sex(M/F) | 21/11 | 24/12 | 20/8 | 16/7 | 10/3 | 0.841<sup>a</sup> |
| Age (y) | 51.2±7.2 | 48.1±9.9 | 49.1±8.6 | 47.4±9.4 | 49.1±9.3 | 0.272<sup>b</sup> |
| Education (y) | 12.7±2.6 | 11.7±2.8 | 12.4±3.0 | 12.2±2.5 | 12.6±3.9 | 0.436<sup>b</sup> |
| NCT (s) | 40.4±10.2 | 56.5±32.1 | 59.8±22.8 | 44.5±21.8 | 45.0±15.6 | 0.009<sup>c</sup>/0.007<sup>d</sup><br>0.357<sup>e</sup>/0.253<sup>f</sup> |
| DST (score) | 47.8±10.5 | 39.2±13.7 | 34.9±16.5 | 48.3±15.0 | 42.3±17.4 | 0.010<sup>c</sup>/0.006<sup>d</sup><br>0.897<sup>e</sup>/0.200<sup>f</sup> |
| Prothrombin time (s) | - | 17.8±6.3 | 17.4±4.5 | 14.8±7.7 | 13.3±6.4 | 0.105<sup>g</sup>/0.022<sup>h</sup> |
| Albumin (mg/dl) | - | 33.2±8.3 | 31.9±5.8 | 40.5±6.0 | 39.3±5.5 | 0.000<sup>g</sup>/0.001<sup>h</sup> |
| Total bilirubin (mg/dl) | - | 78.2±104.0 | 64.9±54.5 | 16.3±10.5 | 15.1±8.1 | 0.001<sup>g</sup>/0.000<sup>h</sup> |
| Blood ammonia (μmol/L) | - | 61.8±21.9 | 84.60±29.5 | 37.0±10.0 | 48.2±22.4 | 0.000<sup>g</sup>/0.000<sup>h</sup> |
| Child-Pugh (A/B/C) | - | 5/14/17 | 0/6/22 | 4/9/10 | 0/2/11 | - |

Data are presented as mean±SD. HCs, healthy controls; OHE, patients with overt hepatic encephalopathy; no-OHE, patients without OHE; NCT, number connection test-A; DST, digit-symbol test;
<sup>a</sup> The p-value was obtained by the Pearson $\chi^2$ test (two-tailed);
<sup>b</sup> The p-value was obtained by the one-way ANOVA test among the three groups (two-tailed);
<sup>c, d</sup> The p-value was obtained by the two-sample t test between pre-LT in no-OHE/OHE group and HCs (two-tailed);
<sup>e, f</sup> The p-value was obtained by the two-sample t test between post-LT in no-OHE/OHE group and HCs (two-tailed)
<sup>g, h</sup> The p-value was obtained by the two-sample t test between post-LT and pre-LT in no-OHE/OHE group (two-tailed);

### 2. Behavior Score Comparison

As expected, OHE and no-OHE groups before LT underperformed compared with HCs - patients took longer to complete NCT tasks (p’s < 0.009) and had lower scores in DST tasks (p’s < 0.01). One month after LT, in comparison to HCs, both the OHE and no-OHE patients showed comparable performance in both DST (p’s > 0.25) and NCT (p’s > 0.20) tasks. The increased performance in DST scores and decreased performance in NCT scores suggested an enhancement of cognitive capability. Also, they may indicate a cognitive recovery after LT for cirrhotic patients regardless of the history of OHE (Table 1).

### 3. Disrupted functional connectivity in cirrhosis

For patients without OHE, using the cluster defining threshold of p < 1×10−4 (explained in Methods), a single network of 17 connections between 16 brain regions was revealed to show decreased functional connectivity in the No-OHE group (p = .037, corrected). We found that the decreased connectivity was mostly found in the connections among sub-cortical nuclei such as Amygdala, Putamen, Pallidum and orbital fontal cortex, as well as insula and cingulate cortex (Figure 2A).

For OHE patients, a single network of 65 connections linking 47 brain regions was revealed with decreased functional connectivity in the OHE group (p = .006, corrected). The connectivity was mostly found in the connections involving the following structures: sub-cortical nuclei such (Amygdata, Putamen, Pallidum), limbic system (temporal cortex, cingulate cortex, parahipocampal gyrus), orbital frontal cortex and other frontal, parietal cortex and cerebellum brain regions (Figure 2B).

NBS was then applied to compare the FC differences between controls and post-LT cirrhosis, and there were no significantly different connections found in both OHE and non-OHE patients.

### 4. Reversed functional connectivity after LT

After LT, using the cluster defining threshold of p < 1×10−4, the disrupted networks found in OHE and non-OHE (as in Fig 2) showed no difference compared to controls (p>0.1).

Furthermore, using the cluster defining threshold of p < 1×10−4, an extra-single network of 19 connections between 17 brain regions was revealed to show increased functional connectivity in the OHE group (p = .009, corrected) after LT relative to the connectivity found before LT. We found that that this increased connectivity was not involved in the disrupted network seen before LT but mostly found in the connections between cerebellum sub-regions, and between cerebellum and occipital cortex, orbital-frontal cortex and limbic systems (Figure 3A). A post-hoc analysis using two-sample t-test revealed that the network strength of this network in OHE post-LT scans was significantly higher than healthy controls (p = 0.008; Figure 3B), suggesting a LT-induced hyperconnectivity for OHE patients only.

### 5. Brain-behavior Model Establishment in healthy controls – Cross-group Validation

A leave-one-out cross-validation was applied to evaluate whether the networks selected from CPM could be generalized to predict unseen subjects. This step was critical because the CPM utilized was constructed using healthy individuals. Once established, the selected functional connectivity would be used as biomarker to evaluate the relationships between functional connectivity and behavioral outcomes after LT for patients’ group. To this end, positive and negative network models described in the method section were used.

In healthy controls, our result showed that the CPM models trained on positive networks predicted the DST task scores of unseen individuals (correlation between predicted and observed DST scores: r = 0.501, p = 0.005), and the models trained on negative networks predicted NCT scores (correlation between predicted and observed, r = 0.446, p = 0.012; Figure 4). Therefore, the positive network model predicting DST scores and the negative network model predicting NCT scores were further utilized to explore the brain-behavior relations in OHE and non-OHE patients.

Because the connections selected in each leave-one-out cycle were not completely identical, we extracted the connections that consistently predicted cognitive test performance in all cycles. In this manner, six connections from the positive network predicting DST and twelve connections from negative network predicting NCT were selected (Figure 5B). The DST and NCT represented similar cognitive functions and we found that their values significantly correlated each other (r = - 0.781, p<0.0001). Thus, we constructed the network that combined connections appearing both NCT negative and DST positive networks, and we called this network “psychomotor network” (PMN). The PMN was then used to predict cognitive performance on patients before and after LT.

**Figure 5.**
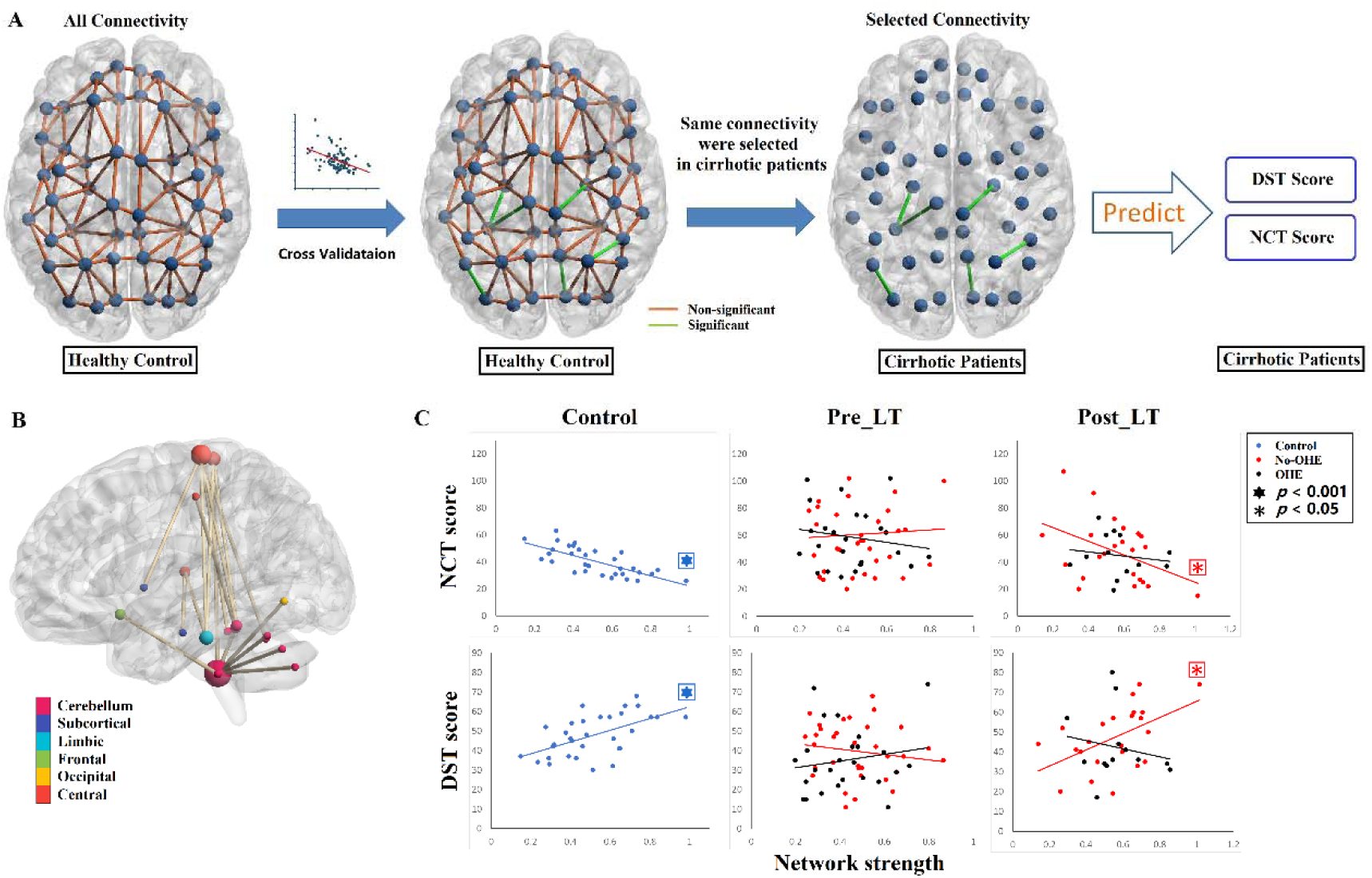
The connectivity-based neuromarkers obtained using connectome-behavior predictive modeling. (A) Connectome-behavior predictive modeling found every single connection that had a meaningful relationship (p < 0.01) to NCT/DST scores in HCs using cross-validation. This meaningful connectivity was then utilized to predict NCT/DST performance for patients before and after the LT. (B) The region pairs with functional connections negatively predicting NCT scores and positively predicting DST scores in controls. These connections formed a single connected network with 19 nodes and 18 connections, and we call this network the psychomotor network (PMN). The predictive connectivity in PMN was mostly found in the connections linked to cerebellum sub-regions, and paracentral cortex. (C) Post-hoc analysis of comparing the predictive power of PMN network strength for NCT and DST scores in both pre-LT and post-LT scans. Before LT, network strength in PMN did not predict NCT/DST scores for both OHE and non-OHE patients. After LT, however, the cognitive test scores were predicted by PMN network strength when analyzing non-OHE patients only (DST: R = 0.517, p = 0.011; NCT: r = −0.424, p = 0.049), but not OHE patients. Non-OHE=none overt hepatic encephalopathy, OHE=overt hepatic encephalopathy, LT=liver transplantation, NCT=number connection test, DST=digit symbol test.

### 6. Brain-behavior relationships in patients before and after LT

For each patient, we first computed network strength by averaging the edge strengths within the PMN, and then conducted linear models relating the PMN network strength to NCT and DST scores. The PMN strength did not predict cognitive performance for pre-LT scans of either OHE or non-OHE patients (p’s > 0.319). However, after LT, the cognitive performance scores in non-OHE were predicted by PMN strength when analyzing their post-LT scans (DST: F(1, 21) = 7.67, p = 0.011, r=0.517, beta=12.58; NCT: F(1, 21) = 4.174, p = 0.049, r=-0.424, beta=-21.33; Figure 5), but the scores in OHE patients were not (p’s > 0.354).

### 7. OHE impacts the neural-behavior relationship after LT

Previously, we found that the PMN strength significantly predicted cognitive performance only in healthy controls and non-OHE patients after LT. Accordingly, we further assessed whether OHE played a moderative role in the predictive relationship between resting state connectivity and cognitive performance in post-LT scans. Moderated regression analyses assessed PMN strength’s association with DST and NCT scores, including OHE as a moderator. These analyses used unstandardized regression coefficients and 95% bias-corrected confidence intervals (CIs) from 10,000 bootstrap estimates (Hayes, 2013; model 3). Results revealed a significant interaction between OHE and non-OHE regarding the relationship between resting state network strength and behavioral performance (p = .029; 95% CI [1.9423, 23.2147]). On the other hand, no such relationships between OHE and non-OHE groups were found before LT.

Similarly, moderated mediation analyses also assessed the association of network strength with NCT scores, including OHE as a moderator. Results revealed a trend of the interaction effect between OHE and non-OHE (p = .061) in post-LT scans. No effect was found in pre-LT scans.

## Discussion

In this study, we investigated the impairment of functional connectivity in cirrhotic patients and the recovery of the impaired functional connectivity after LT by strategically employing whole-brain functional connectivity comparisons and connectome-behavior predictive modeling approaches. We found that the transplant surgery facilitated a recovery of the disrupted functional connectivity in cirrhotic patients. Furthermore, it was shown that OHE patients became abnormally hyperactive in a specific functional network after LT, which was not the part of the initially disrupted network, suggesting the reorganization of the brain network for OHE patients in response to the surgery. Exploring functional connectivity in relation to the psychomotor behavioral performance, we observed that the predictive relationship between functional connectivity and behavioral performance seen in controls was impaired in cirrhotic patients. This impaired relationship was recovered after LT for patients without OHE history only. The moderator role of OHE suggested that the presence of OHE may deform the normal brain connectivity recovery after LT in cirrhotic patients.

### Neural and cognitive recovery difference between OHE and non-OHE patients after LT

Several interesting findings were obtained from the NBS based comparison and post-hoc analysis among groups: 1) OHE patients exhibited larger number of and more widely distributed disrupted FCs than non-OHE patients, suggesting a more serious brain impairment by OHE; 2) No significantly decreased FCs were found in post-LT scans in comparison to controls, i.e. disrupted networks for OHE and non-OHE both recovered; 3) Hyperconnectivity was found in post-LT scans for OHE only, indicating the most significantly increased connections after LT were actually abnormal (higher than in control). Note that both OHE and non-OHE patient didn’t exhibit significantly disrupted connectivity after LT compared to controls. However, the NBS recognized the significantly increased connectivity in OHE patients only after LT compared to before LT. It is not clear that the recovery of the initially disrupted functional connectivity occurs partially or fully (for both OHE and non-OHE patients) because NBS is a relatively rigorous statistical approach and may not capture the difference of the connectivity between pre- and post-op scans. The enhanced functional connectivity identified in OHE patients was unlikely the naturally recovered connectivity as it was not the part of the network that was initially disrupted in the patients before LT. Rather, it could be abnormally established or reorganized network in response to LT under the severe impairment of the brain due to the presence of OHE.

Because hyperconnectivity in response to brain injury or insult has become common in clinical brain network studies (Hillary et al., 2015; Jones et al., 2016; Kim et al., 2020), it might be inappropriate to simply consider the hyperconnectivity after LT as a means of “bigger” recovery for OHE. Instead, hyperconnectivity might represent a reorganization of the functional network in response to the initial damage (probably irreversible) of OHE or another source of brain insults as an adverse surgical effect (Filippini et al., 2009; Hillary & Grafman, 2017). In other words, OHE might induce adaptive and deformed neuroplastic changes or functional re-organization during the brain recovery process following LT. However, it remains to investigate the persistence of the observed hyperconnectivity and its long-term consequences, given chronic engagement of additional neural resources (Hillary & Grafman, 2017).

Psychomotor neurocognitive tests revealed impaired behavioral performance for cirrhotic patients compared to healthy individuals, and performance improvement after LT. Additionally, our investigation of brain-behavior predictions for cirrhotic patients sheds light on the mechanism of cognitive recovery using a generalized psychomotor network (PMN) model. For example, the PMN was not a good predictor of performance before the LT surgery for cirrhotic patients regardless of OHE. After LT, the PMN became significantly associated with cognitive performance for patients without OHE, but not for OHE. Given that the OHE patients also showed better behavioral performance after LT while the association between PMN and cognitive performance was impaired, it was possible that their post-LT behavioral performance was driven by the newly established hyperconnectivity or the network reorganization induced by the possibly severe and irreversible damage of OHE. We further found that one of the key regions in PMN— cerebellum, was largely involved in the hyperconnected network for OHE after LT, suggesting that the hyper cerebellar network established in OHE patients after LT may dissociate the relationship between the PMN and behavioral performance that was observed in healthy individuals. On the other hand, the hyperconnectivity after LT was not observed for non-OHE patients possibly because the damage on the brain due to non-OHE cirrhosis was milder. Accompanied by the recovery of functional connectivity after LT, the PMN could therefore be recovered to drive the behavioral performance for non-OHE patients.

### Functional decoding of psychomotor network (PMN)

Psychomotor ability refers to a wide range of actions involving physical movement related to conscious cognitive processing (Ackerman & Cianciolo, 2000; Kahol et al., 2008). It measures the coordination of multiple cognitive abilities, e.g. attention, visual, executive function, with motor movement. The main hubs (degree >= 5 within PMN) found in the PMN includes paracentral cortex and posterior cerebellum (cerebellum lobe X). Paracentral cortex is a well-known sensorimotor cortex. The cerebellum is mainly involved in the control and coordication of motor movement through multiple mechanism, e.g. timing, spatial evaluation, and sensory acquisition (Stoodley & Schmahmann, 2010). Thus, paracentral cortex and cerebellum in PMN are indicative of motor regulation, which is required in psychomotor tasks. In addition, we also found two less degreed hubs (degree >= 2 within PMN) such as the inferior temporal cortex and the orbitofrontal cortex. The inferior temporal cortex has been reported as an important region associated with visual functions (Woloszyn & Sheinberg, 2012), in particular, playing an important role in the visual-motor coordination. Activities in the orbitofrontal cortex are associated with some higher order cognitive functions like learning (Schoenbaum & Roesch, 2005), rewarding (O’Doherty, Kringelbach, Rolls, Hornak, & Andrews, 2001) and motivation (Arana et al., 2003). In the PMN, orbitofrontal cortex may play roles in the coordination of motor and higher order cognitive functions such as leaning and executive functions.

Note that recent anatomical and functional studies also demonstrated that the cerebellum is involved in a broad range of cognitive functions, besides its historically well-known association with sensorimotor control (King, Hernandez-Castillo, Poldrack, Ivry, & Diedrichsen, 2019; Strick, Dum, & Fiez, 2009). For example, the cerebellum is involved in the executive function (Koziol, Budding, & Chidekel, 2012), attention (Osaka et al., 2004), and emotion process (Guell, Gabrieli, & Schmahmann, 2018). Especially, the cerebellum lobe X, which is one of the main hubs in the PMN, has been suggested as a non-motor functional area in several recent studies (Guell & Schmahmann, 2020; Guell, Schmahmann, Gabrieli, & Ghosh, 2018). It is rather considered associated with visual working memory and visual recognition (King et al., 2019). The clear role of cerebellum in the recovery of OHE still needs to be further addressed in future studies.

### Data-driven approach may find novel and robust neuromarkers

The data-driven approach based on NBS and CPM employed in this study identified novel connections that were used for group comparison and predicting cognitive test outcomes. Previously, hypothesis-driven approaches (García-García et al., 2018; Qi et al., 2012) have limited the selection of the region of interest to the brain regions that are known to be associated with the given cognitive task. Data-driven feature selection methods (M. D. Rosenberg et al., 2016; Zalesky et al., 2010) like the approach adopted in the current study do not have such a limitation. In this regard, our approach found that the neural mechanisms for the processing of cognitive tasks in healthy subjects become impaired in cirrhotic patients and can be recovered after LT in no presence of OHE. This finding implies that specific, but novel brain connections might be involved in cognitive tasks of interest for the patients with cirrhosis. This is more frequently encountered in cognitive tasks that are more taxing e.g. tasks that demand multiple cognitive resources like attention, working memory, motor, and visual spatial components, like psychomotor neurocognitive tasks.

### Limitation and future directions

There are several limitations in current study. The cerebellum plays an important role in fMRI studies for OHE and cirrhosis. Although the recent literature has reported that the cerebellum facilitates successful psychomotor task performance (Medina, Nagel, & Tapert, 2010; Shiroma et al., 2016), the clear role of cerebellum in the recovery of OHE still needs to be further addressed. The current investigation of the brain functional recovery process is conducted based on patients one month after LT. To answer whether the recovery of cognitive function, the recovery of connectivity, the observed hyperconnectivity or functional reorganization in cognitive recovery is temporary or persistent, an extended study that includes longitudinal data with a longer follow-up after LT is needed. Our data-driven method also involves its own problems: it is difficult to clarify the role of the identified functional network or brain regions when their original function is not associated with the target cognitive function.

## Notes

**Conflict of interests** None.

### Competing Interest Statement

The authors have declared no competing interest.

## Reference

Ackerman, P. L., & Cianciolo, A. T. J. J. o. E. P. A. (2000). Cognitive, perceptual-speed, and psychomotor determinants of individual differences during skill acquisition. 6(4), 259.

Ahluwalia, V., Wade, J. B., White, M. B., Gilles, H. S., Heuman, D. M., Fuchs, M., Ganapathy, D. J. L. T. (2016). Liver transplantation significantly improves global functioning and cerebral processing. 22(10), 1379–1390.

Amey, R. C., Leitner, J. B., Liu, M., & Forbes, C. E. (2018). Neural Mechanisms Associated with Semantic and Basic Self-Oriented Memory Processes Interact to Modulate Self-Esteem. bioRxiv, 350926.

Arana, F. S., Parkinson, J. A., Hinton, E., Holland, A. J., Owen, A. M., & Roberts, A. C. (2003). Dissociable contributions of the human amygdala and orbitofrontal cortex to incentive motivation and goal selection. J Neurosci, 23(29), 9632–9638.

Bajaj, J. S., Heuman, D. M., Wade, J. B., Gibson, D. P., Saeian, K., Wegelin, J. A., Stravitz, R. T. J. G. (2011). Rifaximin improves driving simulator performance in a randomized trial of patients with minimal hepatic encephalopathy. 140(2), 478-487. e471.

Bajaj, J. S., Schubert, C. M., Heuman, D. M., Wade, J. B., Gibson, D. P., Topaz, A., Sterling, R. K. J. G. (2010). Persistence of cognitive impairment after resolution of overt hepatic encephalopathy. 138(7), 2332–2340.

Bajaj, J. S., Wade, J. B., & Sanyal, A. J. J. H. (2009). Spectrum of neurocognitive impairment in cirrhosis: implications for the assessment of hepatic encephalopathy. 50(6), 2014–2021.

Barch, D. M., Burgess, G. C., Harms, M. P., Petersen, S. E., Schlaggar, B. L., Corbetta, M., Feldt, C. J. N. (2013). Function in the human connectome: task-fMRI and individual differences in behavior. 80, 169–189.

Bernier, R. A., Roy, A., Venkatesan, U. M., Grossner, E. C., Brenner, E. K., & Hillary, F. G. J. F. i. n. (2017). Corrigendum: Dedifferentiation does not account for hyperconnectivity after traumatic brain injury. 8, 674.

Bharath, R. D., Munivenkatappa, A., Gohel, S., Panda, R., Saini, J., Rajeswaran, J., Biswal, B. B. J. F. i. h. n. (2015). Recovery of resting brain connectivity ensuing mild traumatic brain injury. 9, 513.

Biswal, B., Yetkin, F. Z., Haughton, V. M., & Hyde, J. S. (1995). Functional connectivity in the motor cortex of resting human brain using echo-planar MRI. Magn Reson Med, 34(4), 537–541.

Campagna, F., Montagnese, S., Schiff, S., Biancardi, A., Mapelli, D., Angeli, P., Gatta, A. J. L. T. (2014). Cognitive impairment and electroencephalographic alterations before and after liver transplantation: what is reversible?, 20(8), 977–986.

Cárdenas, A., & Ginès, P. J. C. o. i. c. c. (2011). Acute-on-chronic liver failure: the kidneys. 17(2), 184–189.

Castellanos, N. P., Paúl, N., Ordóñez, V. E., Demuynck, O., Bajo, R., Campo, P., Maestu, F. J. B. (2010). Reorganization of functional connectivity as a correlate of cognitive recovery in acquired brain injury. 133(8), 2365–2381.

Chen, H.-J., Zhang, L., Jiang, L.-F., Chen, Q.-F., Li, J., & Shi, H.-B. J. M. b. d. (2016). Identifying minimal hepatic encephalopathy in cirrhotic patients by measuring spontaneous brain activity. 31(4), 761–769.

Cheng, H., Linhares, B. M., Yu, W., Cardenas, M. G., Ai, Y., Jiang, W., & Cierpicki, T. (2018). Identification of thiourea-based inhibitors of the B-cell lymphoma 6 BTB domain via NMR-based fragment screening and computer-aided drug design. Journal of medicinal chemistry, 61(17), 7573–7588.

Cheng, Y., Huang, L.-X., Zhang, L., Ma, M., Xie, S.-S., Ji, Q., Ni, H.-Y. J. K. j. o. r. (2017). Longitudinal intrinsic brain activity changes in cirrhotic patients before and one month after liver transplantation. 18(2), 370–377.

Cheng, Y., Huang, L., Zhang, X., Zhong, J., Ji, Q., Xie, S., Shen, W. J. M. b. d. (2015). Liver transplantation nearly normalizes brain spontaneous activity and cognitive function at 1 month: a resting-state functional MRI study. 30(4), 979–988.

Eldawud, R., Reitzig, M., Opitz, J., Rojansakul, Y., Jiang, W., Nangia, S., & Dinu, C. Z. (2016). Combinatorial approaches to evaluate nanodiamond uptake and induced cellular fate. Nanotechnology, 27(8), 085107.

Ferenci, P., Lockwood, A., Mullen, K., Tarter, R., Weissenborn, K., & Blei, A. T. J. H. (2002). Hepatic encephalopathy—definition, nomenclature, diagnosis, and quantification: final report of the working party at the 11th World Congresses of Gastroenterology, Vienna, 1998. 35(3), 716–721.

Filippini, N., MacIntosh, B. J., Hough, M. G., Goodwin, G. M., Frisoni, G. B., Smith, S. M., Mackay, C. E. J. P. o. t. N. A. o. S. (2009). Distinct patterns of brain activity in young carriers of the APOE-ε4 allele. 106(17), 7209–7214.

Finn, E. S., Shen, X., Scheinost, D., Rosenberg, M. D., Huang, J., Chun, M. M., Constable, R. T. J. N. n. (2015). Functional connectome fingerprinting: identifying individuals using patterns of brain connectivity. 18(11), 1664.

Forbes, C. E., Amey, R., Magerman, A. B., Duran, K., & Liu, M. (2018). Stereotype-based stressors facilitate emotional memory neural network connectivity and encoding of negative information to degrade math self-perceptions among women. Social cognitive and affective neuroscience, 13(7), 719–740.

García-García, R., Cruz-Gómez, Á. J., Urios, A., Mangas-Losada, A., Forn, C., Escudero-García, D.,. . Giner-Durán, R. J. S. r. (2018). Learning and Memory Impairments in Patients with Minimal Hepatic Encephalopathy are Associated with Structural and Functional Connectivity Alterations in Hippocampus. 8(1), 9664.

Grefkes, C., & Ward, N. S. J. T. N. (2014). Cortical reorganization after stroke: how much and how functional?, 20(1), 56–70.

Guell, X., Gabrieli, J. D., & Schmahmann, J. D. J. N. (2018). Triple representation of language, working memory, social and emotion processing in the cerebellum: convergent evidence from task and seed-based resting-state fMRI analyses in a single large cohort. 172, 437–449.

Guell, X., & Schmahmann, J. (2020). Cerebellar functional anatomy: a didactic summary based on human fMRI evidence. In: Springer.

Guell, X., Schmahmann, J. D., Gabrieli, J. D., & Ghosh, S. S. J. E. (2018). Functional gradients of the cerebellum. 7, e36652.

He, X., Zhu, Y., Lin, Y. C., Li, M., Du, J., Dong, H., & Zhang, L. (2019). PRMT1-mediated FLT3 arginine methylation promotes maintenance of FLT3-ITD+ acute myeloid leukemia. blood, 134(6), 548–560.

Hillary, F. G., & Grafman, J. H. J. T. i. c. s. (2017). Injured brains and adaptive networks: the benefits and costs of hyperconnectivity. 21(5), 385–401.

Hillary, F. G., Roman, C. A., Venkatesan, U., Rajtmajer, S. M., Bajo, R., & Castellanos, N. D. J. N. (2015). Hyperconnectivity is a fundamental response to neurological disruption. 29(1), 59.

Jiang, W., Luo, J., & Nangia, S. (2015). Multiscale approach to investigate self-assembly of telodendrimer based nanocarriers for anticancer drug delivery. Langmuir, 31(14), 4270–4280.

Jiang, W., Wang, X., Guo, D., Luo, J., & Nangia, S. (2016). Drug-specific design of telodendrimer architecture for effective doxorubicin encapsulation. The Journal of Physical Chemistry B, 120(36), 9766–9777.

Jones, D. T., Knopman, D. S., Gunter, J. L., Graff-Radford, J., Vemuri, P., Boeve, B. F., Jack Jr, C. R. J. B. (2016). Cascading network failure across the Alzheimer’s disease spectrum. 139(2), 547–562.

Kahol, K., Leyba, M. J., Deka, M., Deka, V., Mayes, S., Smith, M., Panchanathan, S. J. T. A. J. o. S. (2008). Effect of fatigue on psychomotor and cognitive skills. 195(2), 195–204.

Kim, S. Y., Liu, M., Hong, S.-J., Toga, A. W., Barkovich, A. J., Xu, D., & Kim, H. J. C. C. (2020). Disruption and Compensation of Sulcation-based Covariance Networks in Neonatal Brain Growth after Perinatal Injury.

King, M., Hernandez-Castillo, C. R., Poldrack, R. A., Ivry, R. B., & Diedrichsen, J. J. N. n. (2019). Functional boundaries in the human cerebellum revealed by a multi-domain task battery. 22(8), 1371–1378.

Koziol, L. F., Budding, D. E., & Chidekel, D. J. T. C. (2012). From movement to thought: executive function, embodied cognition, and the cerebellum. 11(2), 505–525.

Liu, M., Amey, R. C., & Forbes, C. E. (2017). On the role of situational stressors in the disruption of global neural network stability during problem solving. Journal of cognitive neuroscience, 29(12), 2037–2053.

Liu, M., Amey, R. C., Magerman, A., Scott, M., & Forbes, C. (2020). The role of startle fluctuation and non-response startle reflex in tracking amygdala dynamics. bioRxiv.

Liu, M., Backer, R. A., Amey, R. C., & Forbes, C. E. (2020). How the brain negotiates divergent executive processing demands: Evidence of network reorganization during fleeting brain states. bioRxiv.

Liu, M., Backer, R. A., Amey, R. C., Splan, E. E., Magerman, A., & Forbes, C. E. J. b. (2020). Context matters: Situational stress impedes functional reorganization of intrinsic brain connectivity during problem solving.

Liu, M., Duffy, B. A., Sun, Z., Toga, A. W., Barkovich, A. J., Xu, D., & Kim, H. (2020, April). Deep Learning of Cortical Surface Features Using Graph-Convolution Predicts Neonatal Brain Age and Neurodevelopmental Outcome. In 2020 IEEE 17th International Symposium on Biomedical Imaging (ISBI) (pp 1335–1338). IEEE.

Liu, M., Kuo, C. C., & Chiu, A. W. (2011, August). Statistical threshold for nonlinear granger causality in motor intention analysis. In 2011 Annual International Conference of the IEEE Engineering in Medicine and Biology Society (pp 5036–5039). IEEE.

Liu, M. T., Kuo, C. C., & Chiu, A. W. (2013). Non-linear Granger causality and its frequency decomposition in decoding human upper limb movement intentions. International Journal of Biomedical Engineering and Technology 34, 12(1), 1–25.

Liu, M., Lepage, C., Jeon, S., Flynn, T., Yuan, S., Kim, J., & Kim, H. (2019, April). A Skeleton and Deformation Based Model for Neonatal Pial Surface Reconstruction in Preterm Newborns. In 2019 IEEE 16th International Symposium on Biomedical Imaging (ISBI 2019) (pp 352–355). IEEE.

Liu, M., & Wang, X. (2017). Beyond the ERPs—Startle Response is Better Outlined by Whole Brain and Spectral EEG Features. Journal of Psychiatry and Brain Science, 2(3).

Ma, H., Irudayanathan, F. J., Jiang, W., & Nangia, S. (2015). Simulating Gram-negative bacterial outer membrane: a coarse grain model. The Journal of Physical Chemistry B, 119(46), 14668–14682.

Medina, K. L., Nagel, B. J., & Tapert, S. F. J. P. R. N. (2010). Abnormal cerebellar morphometry in abstinent adolescent marijuana users. 182(2), 152–159.

Miyake, A., Friedman, N. P., Emerson, M. J., Witzki, A. H., Howerter, A., & Wager, T. D. J. C. p. (2000). The unity and diversity of executive functions and their contributions to complex “frontal lobe” tasks: A latent variable analysis. 41(1), 49–100.

O’Doherty, J., Kringelbach, M. L., Rolls, E. T., Hornak, J., & Andrews, C. (2001). Abstract reward and punishment representations in the human orbitofrontal cortex. Nat Neurosci, 4(1), 95–102. doi: 10.1038/82959

Osaka, N., Osaka, M., Kondo, H., Morishita, M., Fukuyama, H., & Shibasaki, H. J. N. (2004). The neural basis of executive function in working memory: an fMRI study based on individual differences. 21(2), 623–631.

Poldrack, R. A., & Yarkoni, T. J. A. r. o. p. (2016). From brain maps to cognitive ontologies: informatics and the search for mental structure. 67, 587–612.

Poldrack, R. A. J. P. o. P. S. (2010). Mapping mental function to brain structure: how can cognitive neuroimaging succeed?, 5(6), 753–761.

Qi, R., Zhang, L., Wu, S., Zhong, J., Zhang, Z., Zhong, Y., Jiao, Q. J. R. (2012). Altered resting-state brain activity at functional MR imaging during the progression of hepatic encephalopathy. 264(1), 187–195.

Qi, R., Zhang, L. J., Luo, S., Ke, J., Kong, X., Xu, Q., Lu, G. M. J. M. (2014). Default mode network functional connectivity: a promising biomarker for diagnosing minimal hepatic encephalopathy: CONSORT-compliant article. 93(27).

Rosenberg, M., Finn, E., Scheinost, D., Constable, R., & Chun, M. J. T. i. c. s. (2017). Characterizing attention with predictive network models. 21(4), 290–302.

Rosenberg, M. D., Finn, E. S., Scheinost, D., Papademetris, X., Shen, X., Constable, R. T., & Chun, M. M. J. N. n. (2016). A neuromarker of sustained attention from whole-brain functional connectivity. 19(1), 165.

Schoenbaum, G., & Roesch, M. (2005). Orbitofrontal cortex, associative learning, and expectancies. Neuron, 47(5), 633–636. doi: 10.1016/j.neuron.2005.07.018

Shen, X., Finn, E. S., Scheinost, D., Rosenberg, M. D., Chun, M. M., Papademetris, X., & Constable, R. T. J. n. p. (2017). Using connectome-based predictive modeling to predict individual behavior from brain connectivity. 12(3), 506.

Shiroma, A., Nishimura, M., Nagamine, H., Miyagi, T., Hokama, Y., Watanabe, T., Ishiuchi, S. J. T. C. (2016). Cerebellar contribution to pattern separation of human hippocampal memory circuits. 15(6), 645–662.

Sotil, E. U., Gottstein, J., Ayala, E., Randolph, C., & Blei, A. T. J. L. T. (2009). Impact of preoperative overt hepatic encephalopathy on neurocognitive function after liver transplantation. 15(2), 184–192.

Stoodley, C. J., & Schmahmann, J. D. J. C. (2010). Evidence for topographic organization in the cerebellum of motor control versus cognitive and affective processing. 46(7), 831–844.

Strick, P. L., Dum, R. P., & Fiez, J. A. J. A. r. o. n. (2009). Cerebellum and nonmotor function. 32, 413–434.

Tzourio-Mazoyer, N., Landeau, B., Papathanassiou, D., Crivello, F., Etard, O., Delcroix, N., Joliot, M. (2002). Automated anatomical labeling of activations in SPM using a macroscopic anatomical parcellation of the MNI MRI single-subject brain. Neuroimage, 15(1), 273–289. doi: 10.1006/nimg.2001.0978

Umapathy, S., Dhiman, R. K., Grover, S., Duseja, A., & Chawla, Y. K. J. T. A. j. o. g. (2014). Persistence of cognitive impairment after resolution of overt hepatic encephalopathy. 109(7), 1011.

Wang, G., Jiang, W., Mo, S., Xie, L., Liao, Q., Hu, L., & Tong, L. (2020). Nonleaching Antibacterial Concept Demonstrated by In Situ Construction of 2D Nanoflakes on Magnesium. Advanced Science, 7(1), 1902089.

Weissenborn, K., Ennen, J. C., Schomerus, H., Rückert, N., & Hecker, H. J. J. o. h. (2001). Neuropsychological characterization of hepatic encephalopathy. 34(5), 768–773.

Weissenborn, K., Giewekemeyer, K., Heidenreich, S., Bokemeyer, M., Berding, G., & Ahl, B. (2005). Attention, memory, and cognitive function in hepatic encephalopathy. Metab Brain Dis, 20(4), 359–367. doi: 10.1007/s11011-005-7919-z

Woloszyn, L., & Sheinberg, D. L. (2012). Effects of long-term visual experience on responses of distinct classes of single units in inferior temporal cortex. Neuron, 74(1), 193–205. doi: 10.1016/j.neuron.2012.01.032

Zalesky, A., Fornito, A., & Bullmore, E. T. J. N. (2010). Network-based statistic: identifying differences in brain networks. 53(4), 1197–1207.

Zhang, G., Cheng, Y., Liu, B. J. B. i., & behavior. (2017). Abnormalities of voxel-based whole-brain functional connectivity patterns predict the progression of hepatic encephalopathy. 11(3), 784–796.

Zhang, G., Cheng, Y., Shen, W., Liu, B., Huang, L., Xie, S. J. B. i., & behavior. (2017). The short-term effect of liver transplantation on the low-frequency fluctuation of brain activity in cirrhotic patients with and without overt hepatic encephalopathy. 11(6), 1849–1861.

